# Reduction of free-roaming cat population requires high-intensity neutering in spatial contiguity to mitigate compensatory effects

**DOI:** 10.1101/2021.12.02.470990

**Authors:** Idit Gunther, Hadas Hawlena, Lior Azriel, Dan Gibor, Olaf Berke, Eyal Klement

## Abstract

When free-roaming in natural areas, the domestic cat (*Felis silvestris catus*) is ranked high among the most destructive alien species. Near human dwellings, it might risk humans, impair sanitation, and suffer from poor welfare. Cats’ popularity as companion animals complicates their population control. Thus, culling is often replaced by a fertility control method called “Trap-Neuter-Return/Release (TNR),” which is considered more humane. Despite the extensive application of TNR, a long-term controlled study was never performed to test its effectiveness. We present a uniquely designed controlled field experiment for examining TNR effectiveness. The study was performed over a twelve-year period, divided into pre-intervention, mixed- and full-intervention phases, and spanned a 20Km^2^ urban area. Trends of cat, intact-female, and kitten counts, cat reproduction, and carcass reports were compared among study phases and areas with different neutering intensities. The cat population increased during the first two study phases and did not decline in highly neutered populations, presumably due to cat immigration. Expansion of high-intensity neutering to the entire city in the full-intervention phase (>70% neutering percentage) reversed cat population growth, reaching an annual ca. 7% reduction. This population reduction was limited by a rebound increase in cat reproduction and longevity. We conclude that cat population management by TNR should be performed in high-intensity, continuously, and in geographic contiguity to enable population reduction. To enhance management effectiveness and mitigate compensatory effects, we recommend further evaluating an integrated strategy that combines TNR with complementary methods (e.g., vital resource regulation, ill cat euthanasia, and adoption).

**Significance Statement:** Though popular companion animals, domestic cats pose numerous problems when free-roaming, including predation of wildlife, hazards to humans, impaired sanitation, and a decrease in their welfare. Thus, managing their populations is essential. The Trap-Neuter-Return method (TNR, capturing, sterilizing, returning/releasing) is widely employed for managing cat populations. However, there is a lack of long-term controlled evidence for its effectiveness. We examined the outcomes of high-intensity TNR by performing a twelve-year controlled field experiment, the largest to date. Neutering over 70% of the cats caused population decline when applied over contiguous areas. However, it was limited by a rebound increase in reproduction and survival. These findings provide a robust quantification of the limitations and the long-term effectiveness of TNR.

## Introduction

As a generalist predator, the domestic cat (*Felis silvestris catus*) has been listed among the world’s 100 worst non-native species (1). The domestic cat is a fast life-history species (i.e., early maturation, small body size, rapid reproduction) (2-4), which has been distributed around the globe mainly as a pet (5). Cats have been known to form non-domiciliary and often human-independent populations known as free-roaming cats, which tend to cause adverse environmental effects (1, 6-11). The most prominent ecological negative impact of cats is on islands. They are considered responsible for at least 14% of global bird, mammal, and reptile extinctions and are the principal threat to almost 8% of critically endangered birds, mammals, and reptiles (11). Moreover, cats also have a significant ecological effect on the mainland due to direct predation, the transmission of diseases to other species, fear-related effects, and the alteration of demographic processes such as source-sink dynamics (6). As a result of their potential to transmit certain zoonotic diseases and injure humans due to aggressive behavior, free-roaming cats constitute a hazard to public health (7, 8). They might also cause a nuisance to humans, mainly by impairing sanitation (9, 10). These conflicts are intensified in the urban setting where cats form large populations, increasing human-cat interaction (12, 13).

The adverse effects caused by free-roaming cats have increased the motivation to artificially manage their populations, aiming either to diminish their related nuisances or to preserve natural ecosystems (14-19). Animal population management is based on two strategies: resource limitation in the habitat or actions applied to individuals. The latter strategy can be further divided into culling and fertility control methods. While culling aims to increase mortality above the natural rate, fertility control aims to decrease reproduction below the natural rate (20). Culling has been successful in eradicating cat populations on certain islands (14, 16, 17). However, there are several examples of the failure of culling to accomplish population control in fast life-history species, such as voles (21), mice, rats, jirds (22), rabbits (23), cats (24, 25), and foxes (26). In contrast, information on the efficacy of fertility control on vertebrate populations is scarce (27). Specifically for free-roaming cats, theoretical studies predicted that culling performs better than Trap-Neuter-Return (TNR), a common method for controlling cat fertility (20, 28-31). However, despite these predictions, the TNR method has been progressively implemented in cat populations over widespread areas instead of culling, mainly due to moral considerations and public opinion (18, 32-36).

A growing collection of studies has examined the effects of TNR on cat population dynamics, however ours is the first controlled study conducted to observe the long-term effectiveness of TNR. Past studies have yielded inconsistent finding with several finding decreases in population growth indicators (37-52), and others finding stabilization or even increases (53-58). These inconsistent results might stem from differences in management duration and efforts, the examined populations (e.g., closed versus open populations, small versus large-scale populations), or the study environment. The inference of the overall long-term consequences of TNR is further limited due to: absence of control (37-41, 43-45, 48-52, 54, 56, 57), combining TNR with other control tactics (i.e., adoption and euthanasia of clinically ill or retrovirus-positive cats) (37-46, 49-52), short-term follow-up (37, 38, 42, 45, 46, 51, 53-56,58), small sample size (37, 38, 40, 43, 45, 47, 49, 50, 52-54, 58), relying on indirect indices of population growth (38,39, 42, 46, 49, 51, 52, 57), and examining populations in secluded areas (41, 44).

Boon et al. (59) called for a long-term and large-scale study in cat populations to close the knowledge gap and overcome shortcomings of previous studies. Here we present the results of a 12-year longitudinal large-scale experiment. We assessed the effect of sterilization (i.e., spaying/neutering) on the long-term temporal and spatial dynamics of free-roaming cat populations. While adjusting for environmental factors, we compared cat population dynamics before and at partial and full implementation of a TNR program. We found that population compensatory mechanisms may limit the magnitude of the neutering effect. Thus, TNR should be performed continuously, at high intensity, and in spatial contiguity to enable long-term cat population regulation.

## Results

### Neutering intensity across statistical areas and study phases

The study was set to assess long-term TNR effectiveness over time and space (**Figures 1, S1**). In the 1^st^ (pre-intervention) phase, neutering was not applied. In the 2^nd^ (mixed-intervention) phase, high-intensity neutering was applied in about half of the city, and in the 3^rd^ (full-intervention) phase, TNR was applied in the entire city. We confirmed the neutering status in 96% of the surveyed cats by ear-mark detection, counting both unneutered (un-marked) females (queens) and males and neutered (marked) cats, along with kittens. Repeated annual surveys were performed during September and October of 2012, 2013, 2014, and 2018. Overall, in the 50 surveyed statistical areas, 13,718 cat-observations were documented during the study period, of which 1,486 were kittens. The phase-and group-specific neutering percentages corroborated the respective TNR intensity (**Table 1, Table S1**). For a visual perspective of the municipal TNR intensity across statistical areas and study phases, see **Figure S2**.

**Figure 1:**
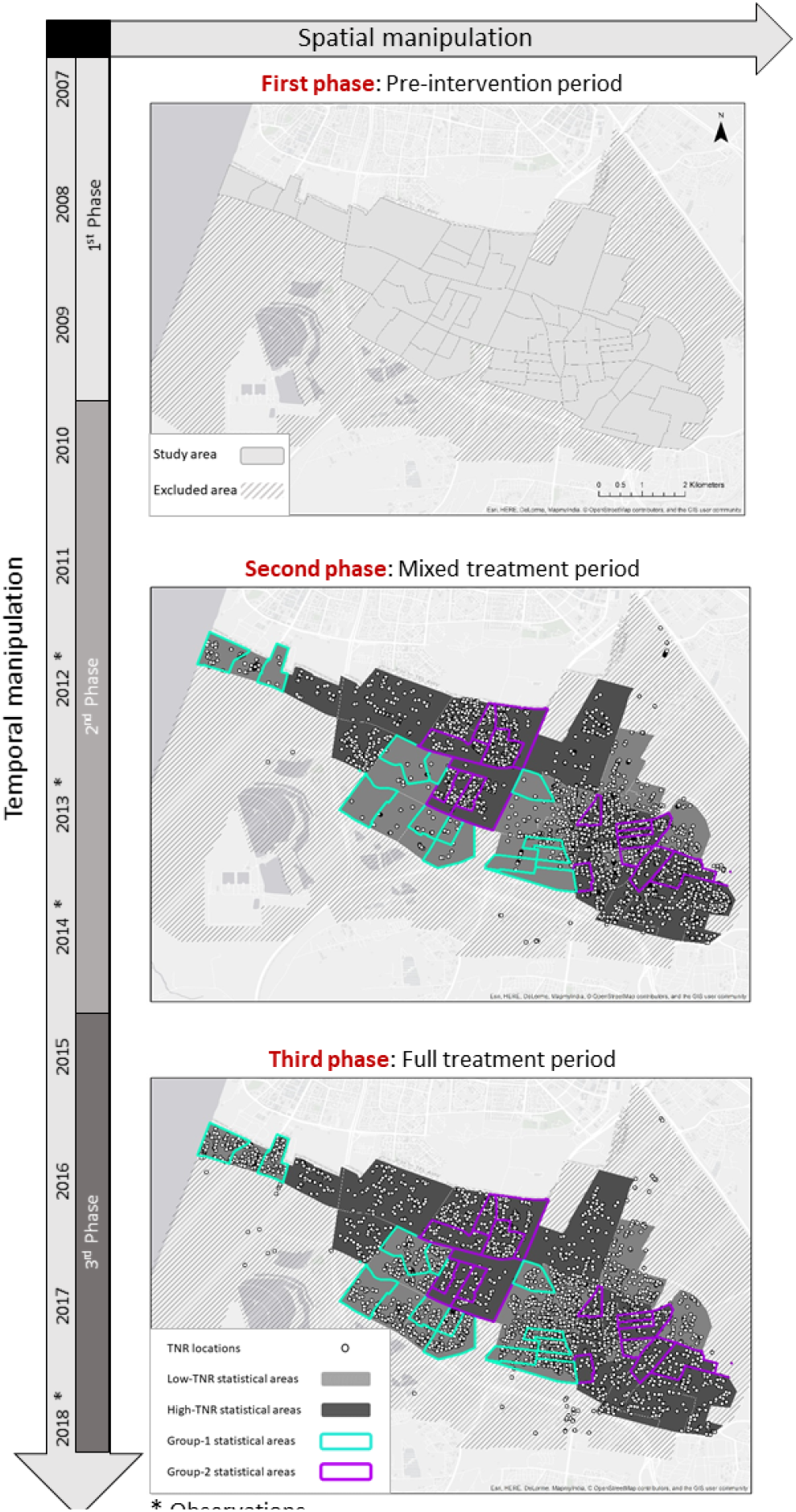
The experimental design was set to assess long-term TNR effectiveness over time and space. The study period was divided into three phases: 1^st^ phase – pre-intervention period -between 2007 to the end of 2009; 2^nd^ phase -mixed-intervention -allocating TNR to half of the city neighborhoods between the end of 2009 to end of 2014; 3^rd^ phase, full-intervention period, allocating TNR to the entire city between the end of 2014 and the end of 2018. Dots present locations of TNR in the city area. The study area was composed of 50 statistical areas (presented as polygons in the upper map; see also Fig. S1), 24 of which are termed high-TNR, as they were treated in both TNR phases, and the other 26, termed low-TNR, as they were mainly treated during the full-intervention phase. From these high-and low-TNR statistical areas, 15 and 12 areas with the highest and lowest neutering percentage at the 2014 survey, respectively, were served as the main study units (Group-1 and -2). Survey years are marked by an asterisk.

**Table 1:**
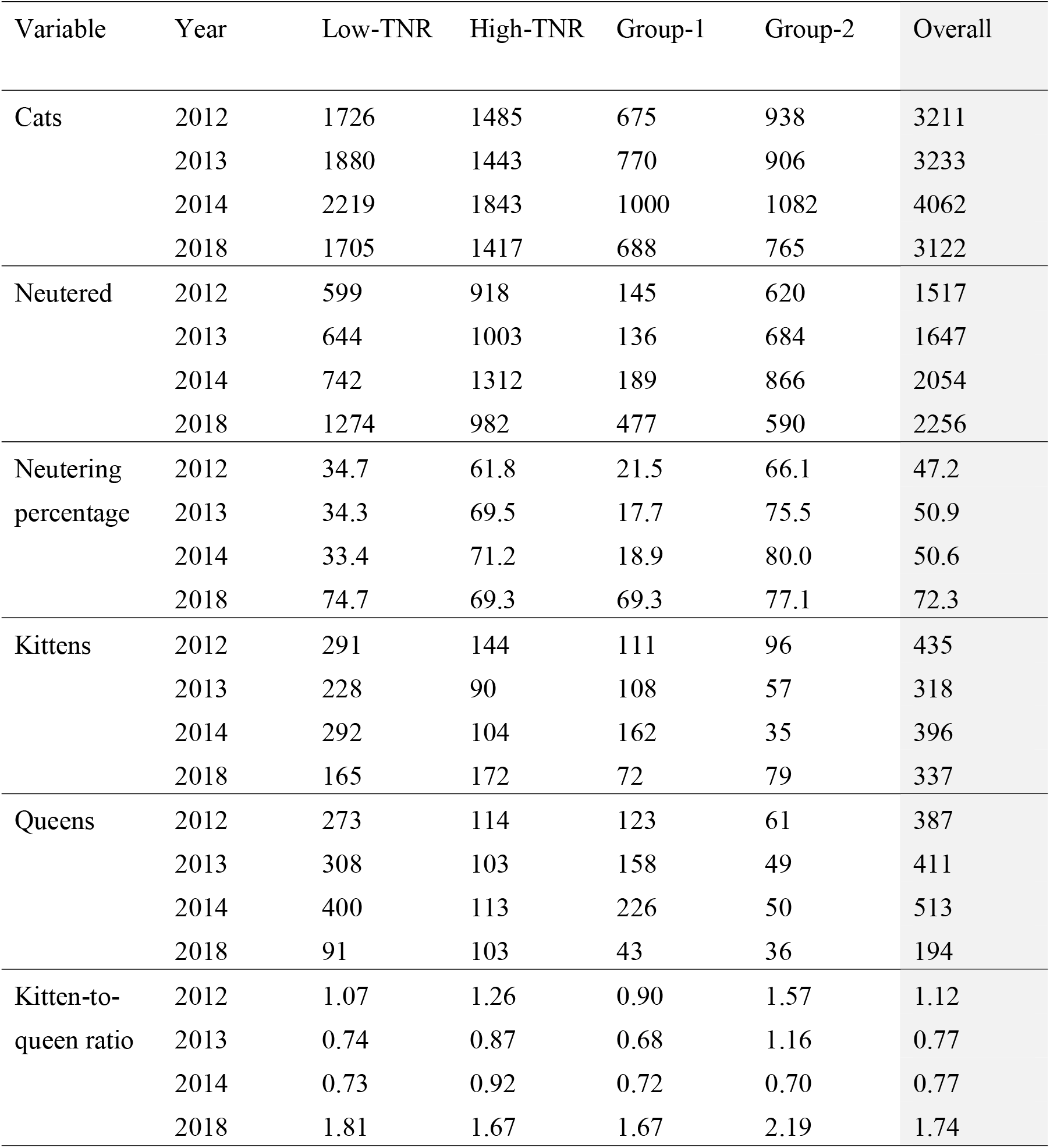
Annual counts of observed cats, neutered cats, neutering percentage, kittens, queens, and kitten-to-queen ratio in the low-TNR (n=26), high-TNR statistical areas (n=24), Group-1 (lowest neutering intensity, n=12) and Group-2 (highest neutering intensity, n=15).

At the end of the mixed-intervention phase (end of 2014), the overall neutering percentages in the entire surveyed area, the low TNR (n=26) and high TNR (n=24) statistical areas, were 51, 33, and 71%, respectively (**Table 1**). To compare the effect of the highest-intensity neutering to the lowest-intensity, we further divided the statistical areas into quartiles according to the observed neutering percentage in that year (2014). Group-1 (n=12) represents the lowest neutering intensity and consists of the statistical areas with neutering percentage at the lowest quartile. Group-2 (n=15) represents the highest neutering intensity and consists of the statistical areas with neutering percentage at the highest quartile (see **Figure S1** for the detailed spatial study design). At the end of the mixed-intervention phase, the overall neutering percentages in Group-1 and Group-2 were 19 and 80%, respectively. At the end of the full-intervention phase (end of 2018), the neutering percentage in the entire area was 72%, and in the low TNR, high TNR, Group-1, and Group-2, the overall percentages were 75, 69, 69, and 77%, respectively (**Table 1, Figure 2A**).

**Figure 2:**
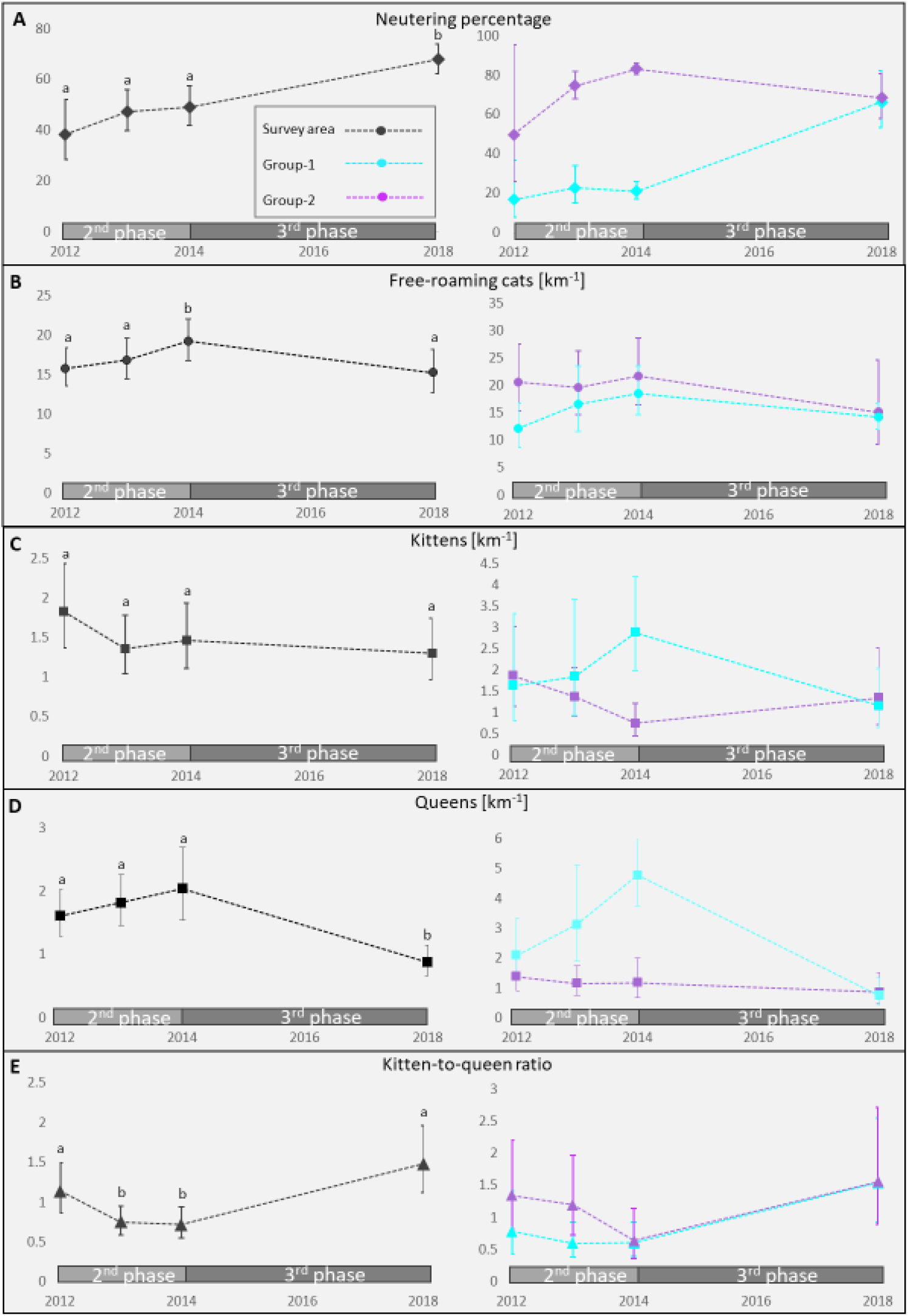
Annual trends (geometric mean±CI_95%_) of (**A**) neutering percentage, (**B**) free-roaming cat counts, (**C**) kitten counts, (**D**) queen counts, and (**E**) kitten-to-queen ratio in the city of Rishon-LeZion (black, n = 50 statistical areas), and in Group-1 (turquoise, lowest neutering intensity, n=12 statistical areas) and Group-2 (purple, highest neutering intensity, n=15 statistical areas). Observations were performed in 2012–2014 and 2018. Superscript letters represent years, where observations differ.

### Environmental human-related factors affecting cat population dynamics

We modeled year-based cross-sectional data (adjusting for neutering percentage) to screen for potential human-related factors that might influence cat population dynamics. As expected, we found a consistent negative association between neutering percentage and kitten counts (**Table S3**). Interestingly, in some years, the cat counts were positively associated with the neutering percentage (i.e., 2012 and 2018), indicating possibly higher TNR efforts in areas with higher, rather than lower, cat densities. In addition, cat population size parameters, cat counts, and resident reports of cat carcasses and reproduction were positively correlated with the human population density (**Table S3**). Though less consistent, the volume of accessed-waste-bins was also positively correlated with some cat population parameters. Therefore, we adjusted for human population density and accessed-waste-bins by including them in the analysis of neutering association with cat population trends. The other tested human-related factors were not consistently associated with the cat population parameters. We did not find a consistent spatial correlation in these models, and therefore it was not further included in the trend models.

### Survey results: Cat and kitten counts across statistical areas and study phases

During the mixed-intervention phase (2012-2014), we found an overall increase of 26.5% in the cat counts in the surveyed area. Therein, we found an average annual growth of 20.7% in Group-1, which was exposed to the lowest neutering intensity (**Table 1, Figure 2B**). At the same time, the cat counts remained stable in Group-2, which was exposed to the highest neutering intensity (**Table 2**). In contrast, during the full-intervention phase, we found an overall reversal of population growth, with a 23.1% reduction in the cat counts in the entire surveyed area. During this phase, the annual decrease in the two treatment groups was similar (ca. -7% annually; **Figure 2B)**.

**Table 2.** Annual trends of the cat and kitten counts, kitten-to-queen ratio (surveyed during the mixed-and full-intervention phases), and cat carcass and reproduction data (received from resident reports during all the three study phases). The trends are depicted for Group-1, (lowest neutering intensity, n=12), and Group-2, (highest neutering intensity, n=15). All the models are adjusted for human population density and accessed waste bins.

| Population parameter | Phase | Measurement period | Annual trend for Group-1 (CI<sub>95%</sub>) [%] | Annual trend for Group-2 (CI<sub>95%</sub>) [%] | P-value |
| --- | --- | --- | --- | --- | --- |
| Cat counts | 2<sup>nd</sup> phase | 2012-2014 | 20.72 (8.19 to 34.71)<sup>a1*</sup> | 4.02 (-5.62 to 14.65)<sup>a2</sup> | 0.046 |
|  | 3<sup>rd</sup> phase | 2014-2018 | -7.95 (-13.09 to -2.50)<sup>b1</sup> | -7.83 (-12.29 to -3.15)<sup>b2</sup> | 0.973 |
| Kitten counts | 2<sup>nd</sup> phase | 2012-2014 | 20.20 (-11.31 to 62.91)<sup>a1</sup> | -42.39 (-58.19 to -20.60)<sup>a2</sup> | 0.001 |
|  | 3<sup>rd</sup> phase | 2014-2018 | -14.14 (-27.00 to 1.00)<sup>b1</sup> | 21.37 (3.20 to 42.74)<sup>b2</sup> | 0.003 |
| Queen counts | 2<sup>nd</sup> phase | 2012-2014 | 40.40 (13.31 to 73.97)<sup>a1</sup> | -11.21 (-30.92 to 14.12)<sup>a2</sup> | 0.007 |
|  | 3<sup>rd</sup> phase | 2014-2018 | -38.34 (-46.47 to -28.98)<sup>b1</sup> | -8.76 (-20.50 to 4.71)<sup>a2</sup> | <0.001 |
| Kitten-to-queen ratio | 2<sup>nd</sup> phase | 2012-2014 | -7.72 (-36.10 to 33.28)<sup>a1</sup> | -28.48 (-50.60 to 3.55)<sup>a2</sup> | 0.338 |
|  | 3<sup>rd</sup> phase | 2014-2018 | 29.88 (7.40 to 57.08)<sup>a1</sup> | 23.35 (2.26 to 48.28)<sup>b2</sup> | 0.706 |
| Carcass reports | 1<sup>st</sup> phase | 2007-2009 | 7.96 (-1.44 to 18.26)<sup>a1</sup> | 6.01 (-1.62 to 14.23)<sup>a2</sup> | 0.761 |
|  | 2<sup>nd</sup> phase | 2010-2014 | 0.88 (-3.04 to 4.95)<sup>a1</sup> | -12.02 (-15.17 to -8.75)<sup>b2</sup> | <0.001 |
|  | 3<sup>rd</sup> phase | 2015-2018 | -13.59 (-19.02 to -7.79)<sup>b1</sup> | 0.57 (-4.82 to 6.27)<sup>a2</sup> | <0.001 |
| Reproduction reports | 1<sup>st</sup> phase | 2007-2009 | 8.02 (-8.03 to 26.87)<sup>a1</sup> | -5.70 (-18.44 to 9.04)<sup>a2</sup> | 0.219 |
|  | 2<sup>nd</sup> phase | 2010-2014 | 3.89 (-2.95 to 11.21)<sup>a1</sup> | -24.50 (-29.94 to -18.63)<sup>b2</sup> | <0.001 |
|  | 3<sup>rd</sup> phase | 2015-2018 | -7.23 (-17.18 to 3.91)<sup>a1</sup> | 22.56 (9.96 to 36.59)<sup>c2</sup> | <0.001 |

\* The superscript letters represent comparisons between pairs of phases within each group of statistical areas (pairs with insufficient evidence for a difference are labeled with the same superscript letter).

As opposed to cats, we did not find a significant reduction in the kitten counts in the entire surveyed area (**Table 1, Figure 2C**). This overall stability was a result of opposite trends in the two treatment groups. As expected, during the mixed-intervention phase, kitten counts increased annually by 20.2% in Group-1 but decreased by 42.4% in Group-2. Whereas, during the full-intervention phase, these trends reversed, with a decrease of 14.1% in Group-1 and an increase of 21.4% in Group-2 (**Table 2**). The latter increase in kitten counts in Group-2, during the full-intervention phase, was unexpected. A possible explanation for this increase is a reduction in neutering intensity in some statistical areas in this group during the full-intervention phase. To examine this possibility, we reanalyzed the data of both groups, excluding statistical areas in which the neutering percentages at the end of the full-intervention phase were below 70%.

Here, too, we found the same trend reversal in kitten counts (**Table S4**). Therefore, a decrease in neutering intensity in some statistical areas is unlikely to explain the observed increase. An alternative explanation is that reduced agonistic behavior in neutered cats enhanced compensatory mechanisms such as increased kitten survival, a rise in the frequency of pregnancies per queen, or increased litter size due to lower competition. Assuming this hypothesis was correct, we predicted to find an overall increase in the kitten-to-queen ratio in the full-intervention study phase. We indeed found a 2.25-fold increase during this phase (**Table 1, Figure 2E**). This increase was significant in both treatment groups (**Table 2**). In Group-2, it occurred due to a significant increase in the kitten number and stabilization of the queen number. In Group-1, the queen number decreased annually by 38% shortly after introducing high-intensity neutering, while a 14% insignificant decrease in the kitten number was observed (**Table 2, Figure 2C,D,E**). No significant reduction in the kitten-to-queen ratio was observed in Group-2 during the mixed-intervention period, possibly due to parallel immigration of intact females.

### Resident report results: Cat carcasses and reproduction events across statistical areas and study phases

We complemented the survey data analysis with those of resident reports on cat reproduction events and cat carcasses. This dataset increased the study resolution in both space (covering more statistical areas) and time (daily reports from 2007 to 2018). To examine the construct validity of these reports, we tested the correlation of reproduction reports with the surveyed kitten counts and found a strong correlation between the two datasets (Pearson correlation coefficient=0.55, p<0.001, **Figure S6**).

In addition, to ensure that these reports represent trends unique to cats, we compared them with the number of all available reports to the call center (after excluding cat-related reports) and with the reports of all carcasses (excluding cats and including canines, poultry, rodents, hyraxes, hedgehogs, equines, reptiles, boar and tortoises). The trends of these two latter report types showed a consistent increase during the study period (**Figure S4**). Overall, this increased trend suggests that people’s awareness or tendency to use the call center increased with time. Unlike the general trends, the reports of cat carcasses and reproduction were linked to the application of TNR. As with the general reports, both showed a mild positive trend during the pre-intervention phase. However, this was followed by a negative trend, from the onset of the mixed-intervention phase until one year after its end (from 2011 to 2015) (**Figure 3**). The magnitude of this negative trend was ca. 50%, starting at 325 and 120 and reaching a low of 200 and 60 of carcasses and reproduction reports per month, respectively. Notably, during the mixed-intervention phase, the number of carcass and reproduction reports decreased while the cat counts increased (**Table 1, Figure 2**), indicating a reduction in cat reproduction and mortality. One year after the onset of the full-intervention phase (2016-2018), the number of carcass reports stabilized, but the number of reproduction reports showed an unexpected increase (**Figure 3**). This increase suggests a possible rebound effect of TNR on reproduction.

**Figure 3:**
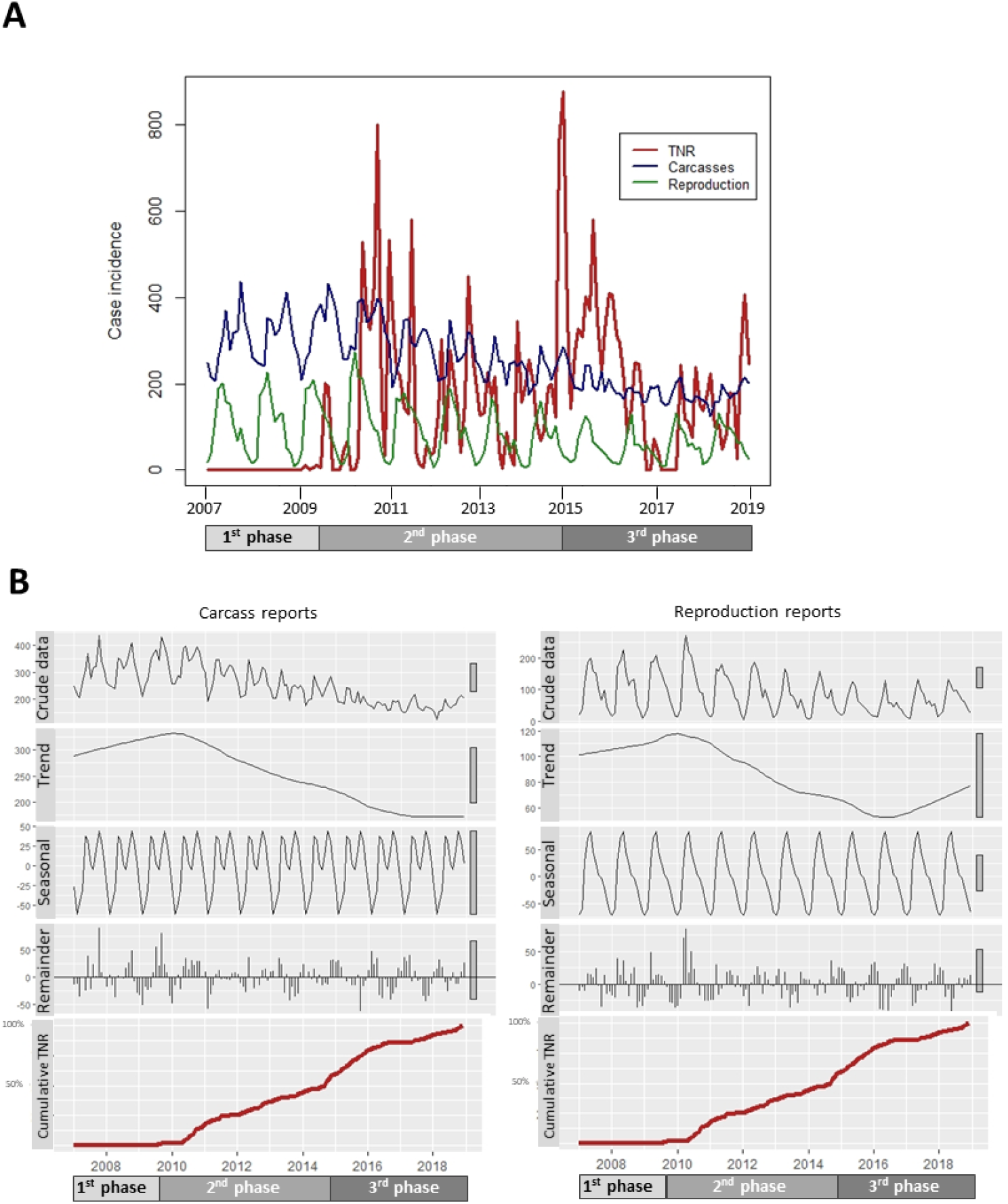
Municipal TNR efforts and time series decomposition plots of the monthly carcass and cat-reproduction citizen reports in the entire city of Rishon-Lezion, between 2007-2018 (the crude data is divided into three compartments: trend, seasonality and remainder). (**A**) Monthly-based time series plots of municipal TNR actions (n = 22,144), cat carcass reports (n = 36,544) and reproduction reports (n = 12,217) from January 2007 to December 2018 across the entire city (61 statistical areas). (**B**) Time series decomposition plots of carcass and reproduction reports linked with cumulative TNR percentage. The 1^st^ phase refers to the pre-intervention period (before the initiation of the TNR program), the 2^nd^ phase refers to the mixed-intervention period (applying TNR in half of the city), and the 3^rd^ phase refers to the full-intervention period (applying TNR in the entire city).

Both report types exhibited a prominent seasonal pattern: carcasses peaked twice, in May-June and in October, whereas reproduction reports peaked only once in April-May (**Figure 3**). Unlike these reports, the general reports to the call center peaked in July-August (**Figure S4**), showing that the seasonal pattern of carcass and reproduction reports is specific for cats and did not represent human behavior tendency to refer to the call center. While the seasonal pattern of cat reproduction reports was maintained throughout the study, the seasonality of carcass reports gradually diminished from the onset of TNR until it disappeared during the full-intervention phase (see ‘crude data’ in **Figure 3b**). Thus, it implies that TNR resulted in a decrease in mortality and a decrease in the seasonal (reproduction-linked) mortality pattern.

We next compared the carcass and reproduction report trends between the treatment groups. Both trends coincided with TNR efforts (**Figure 4, Table 2**). In the pre-intervention phase, the numbers of carcass and reproduction reports were stable. During the mixed-intervention phase, both carcass and reproduction reports declined considerably in Group-2 (−12% annually for carcass and -24.5% for reproduction reports), while their numbers were stable in Group-1 (0.9% for carcass and 3.9% for reproduction reports). The number of carcass reports decreased considerably in Group-1 (−13.6%) in the full-intervention phase, indicating a decrease in cat number and/or mortality. Similar to kitten counts, in Group-2, a considerable rebound increase of reproduction reports occurred during the full-intervention phase (22.6%). We found similar trends when the analysis was repeated after excluding statistical areas in which the neutering percentages at the end of the full-intervention phase were below 70%. (**Figure S5, Table S4**).

**Figure 4:**
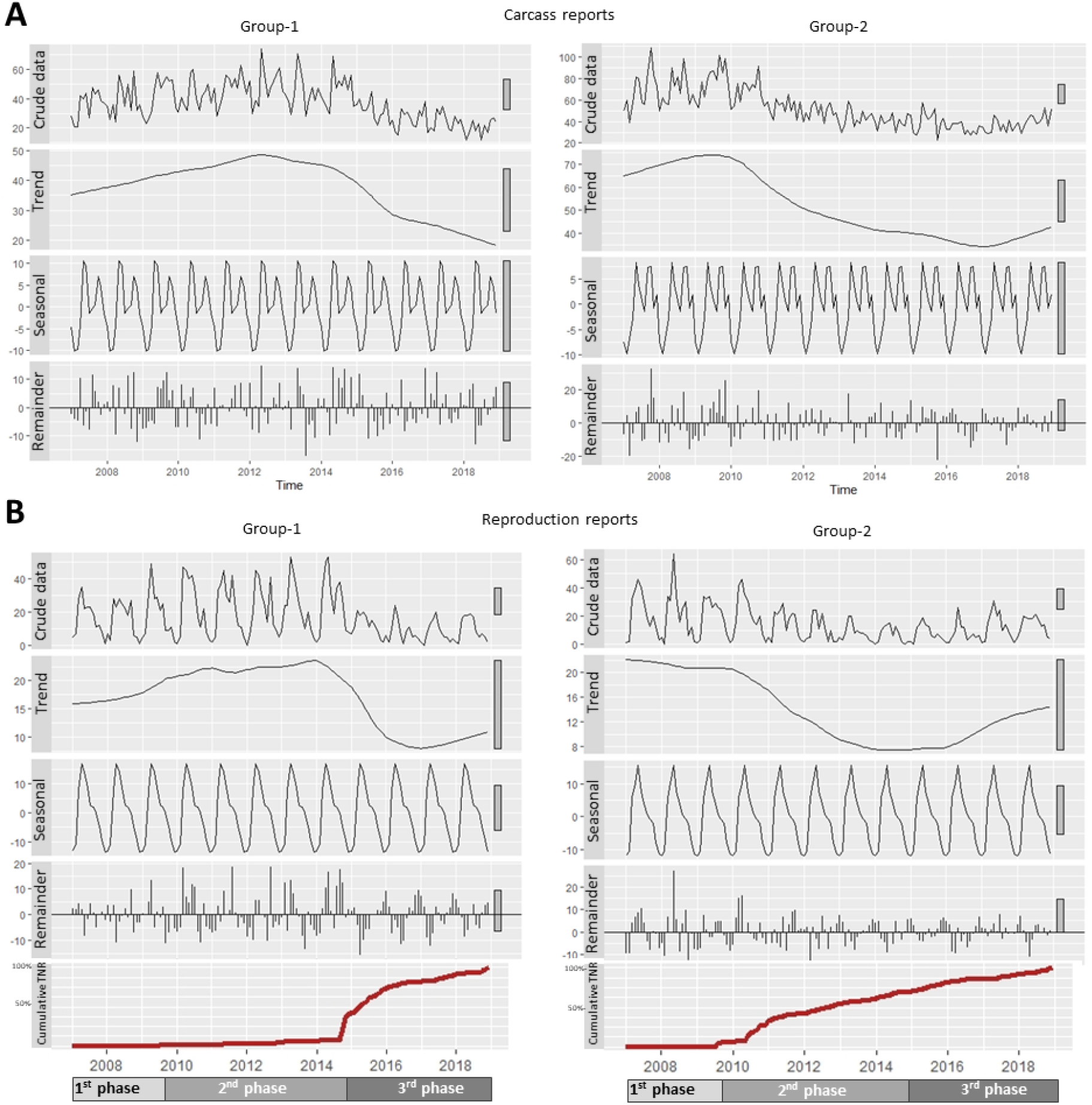
Municipal TNR efforts and time series decomposition plots of the monthly carcass and cat-reproduction citizen reports in group 1 and group 2 statistical areas in the city of Rishon-Lezion, between 2007-2018 (the crude data is divided into three compartments: trend, seasonality and remainder) of (**A**) cat carcass reports and (**B**) cat reproduction reports in Group-1 (lowest neutering intensity, n = 12) compared to Group-2 (highest neutering intensity, n = 15). The 1^st^ phase refers to the pre-intervention period (before the initiation of the TNR program), the 2^nd^ phase refers to the mixed-intervention period (applying TNR primarily in Group-2), and the 3^rd^ phase refers to the full-intervention period (applying TNR in both groups).

## Discussion

The current study is the first long-term controlled field experiment examining TNR effectiveness in free-roaming cat populations. Using a unique experimental design in which we manipulated neutering intensity across time and space, we distinguished between short-term and long-term population consequences of TNR. To achieve high-resolution data we used two independent data-collection methods-a repeated cross-sectional annual survey and daily resident reports. Parameters of population dynamics derived by both methods showed the same trends, suggesting that TNR can reduce population size, but only when applied at high rates in spatial contiguity. Below, we discuss the evidence suggesting processes that might challenge TNR effectiveness and thus should be considered when applying it. These include compensatory effects, such as increased cat survival and/or litter size, as well as possible influx of immigrant cats into vacant niches. We then compare our results to those generated by theoretical simulations, discussing the TNR option compared to its alternatives, and end by offering directions for further simulations and planning strategies.

### Maintaining high neutering intensity in spatial contiguity is required to mitigate counteracting effects

During the mixed-intervention phase, the cat population growth in Group-2 was considerably lower than in Group-1. However, the cat population did not decline in Group-2 despite maintaining a neutering percentage of 80% (**Figures 2A, B**), a number previously suggested as sufficient for reducing the population size of free-roaming cats (28, 60). We propose that during the mixed-intervention phase, the TNR effect on cat number was counteracted by the immigration of intact cats from unneutered surrounding populations. It was previously shown that sterilization-enhanced behavioral changes resulted in increased immigration of free-roaming cats from low-into high-TNR regions (53). This important population compensatory process was also documented in other managed species (reviewed in Ransom et al. 2014 (27)). The study area boundaries may have enabled long-distance cat immigration mainly from the neighboring northern cities, where municipal TNR campaigns were not implemented. However, we believe it was negligible due to the small home range of urban cats (60, 61).

During the full-intervention phase, an average annual reduction of more than 7% was observed in groups 1 & 2. Our surveys further indicated that during the four years of this phase, the cat population in the entire city declined by ca. 25%. It is possible that the cat counts decreased in the entire city due to temporal effects unrelated to TNR. For example, a change in vital resource availability (e.g., decreased feeding or change in sanitary conditions), decreased abandonment, or increased adoption. However, this possibility is unlikely for three reasons: First, the finding of cat counts reduction was adjusted for changes in the environmental factors influencing cat population size (i.e., human population density and the per human accessible waste bins). Second, the survey periods following each intervention change were not longer than 4−5 years. A marked change in human behavior to opposite directions both spatially and temporally in such short periods is unlikely. Third, though abandonment and adoption of free roaming cats do occur in Israel, they constitute only a negligible factor relative to the remarkable density of the free-roaming cats’ population (751-2300/Km^2^ (61, 62)). The most probable alternative explanation for the cat population trend changes between the phases is the expansion of high neutering intensity (>70%) to the entire city during the full-intervention phase. This expansion may have reduced the effect of the immigration of cats from low TNR to high TNR areas. We conclude that maintaining high-intensity neutering in spatial contiguity is a prerequisite for reducing cat population size.

### Limiting processes should be considered in TNR programs

Along with the encouraging cat population decline, our findings support the occurrence of several compensatory effects, limiting TNR effectiveness: **Reduced mortality**; Cat carcass trend reversed from an increase during pre-intervention to a constant decrease during the mixed-intervention phase. This decrease opposed growth in cat population size at this phase. These opposite trends indicate that the reduced carcass reports are due to reduced mortality following TNR. **Reduced reproduction-related mortality**; The diminished seasonal pattern of carcass reports following increased neutering intensity (see ‘crude data’ in Fig 3B and Fig 4b) indicates a decreased reproduction-related mortality. It was previously shown that in the Northern hemisphere, kittens present a seasonal appearance (4, 10, 52, 64) and experience higher morbidity and mortality than adults (53, 63, 64). Moreover, the mortality of intact cats was higher than neutered ones (65, 66), probably due to associated reproductive behavior. **Increased fertility and reduced kitten mortality**; While the neutering percentage exceeded 72% in the entire city during the full-intervention phase, a considerable increase in the kitten-to-queen ratio was documented in both study groups. This increased ratio may stem from several plausible explanations: an elevation in kitten survival, a rise in the frequency of pregnancies per queen, and a rise in litter size. This conclusion is supported by a rise in cat reproduction reports in Group-2 during the full-intervention phase. Increased fertility and decreased mortality of juveniles and adults were previously reported to follow fertility control in domestic cats (34, 53, 63, 64) and other vertebrates (27, 67-71). Both compensatory processes could result from higher food availability and decreased resource competition following population decline (64, 72). In addition, they could be caused by diminished agonistic behavior in neutered male cats (73). These mechanisms, together with cat immigration, might limit the TNR-derived decrease in cat population size.

### Comparison of our findings to the results of cat population simulation models

Several theoretical simulations aimed to predict the long-term consequences of TNR and provide specific management insights (20, 28, 30, 31). Of these, only two studies examined the effect of TNR in an open population (20, 30). In these studies, population reduction was achieved by implementing TNR, though the immigration of non-neutered cats limited its effectiveness. This situation resembles the mixed-intervention phase in the current study when untreated areas surrounded treated areas. Interestingly, in our study, during this phase, the cat population did not decline. There are at least two possible explanations for this discrepancy. First, we assume that similar to other population dynamics, immigration is a density-dependent process. Thus, the immigration rate may depend on density differences of both total cats and intact cats between the managed and surrounding populations (53). By modeling immigration as a density-independent process, the theoretical models might have underestimated this process’s magnitude. Second, these models did not include other density-dependent population compensatory mechanisms such as increased survival and reproduction, which probably occurred in our study.

### Comparison of TNR to culling

When culling or other means of cat removal were applied intensively, they eradicated cat populations on certain islands (14, 16, 17). Although culling may be efficient, concurrent similar compensatory mechanisms to those observed in our study might also limit its effectiveness (24, 25). In some environmental contexts, TNR is considered as an alternative to culling. Since the public acceptance of TNR is better than culling (9, 35, 36, 74), it improves public cooperation with policy-makers, which is an important factor for population reduction success. The merits of TNR go beyond those achieved by population reduction and include improved welfare (64) and a reduction in cat-related nuisances, which are mainly associated with reproduction (38, 75). On the other hand, the costs of TNR implementation should be taken into account. Especially challenging is the maintenance of high-intensity neutering for long periods, considering the significant decrease in trapping efficiency when the percentage of neutered cats is high. More than one million dollars ($US) were invested in the current project over the nine years of TNR implementation. This cost should be compared to alternative population management programs, considering the differences between countries, economies, jurisdictions, and ethical perceptions.

### Complementary approaches that can further improve TNR benefits

Combining TNR with removing a significant portion of the cat population (i.e., euthanasia of ill cats and adoption) can improve its effectiveness (40, 42-44). However, the removal of cats might not be applicable everywhere. Our study’s analysis of the environmental factors influencing cat populations highlights an alternative approach to complement TNR. The results show an association between vital anthropogenic resources (i.e. accessed-waste-bins and human density, which may be associated with higher food availability to cats) and cat population parameters. As suggested by Boone (2015), limiting such resources might be combined with TNR to reduce carrying capacity along with the reduction in population size (76). Despite population reduction, neutered cats might continue to hunt (77, 78). Therefore, TNR is not an effective means by itself for preventing predation. It was shown that manipulating food content (e.g. providing high-meat protein and grain-free diet) and applying environmental enrichment (e.g. object play) reduced cat tendency to hunt (79), and placing collar-mounted devices interfered with hunting (80). Such measures can be therefore combined with TNR to diminish this adverse effect further.

### Strengths and limitations

The current experiment is unique not only in the controlled and long-term design but also in the large number and depth of data of neutered and observed cats, the large covered urban area which is systematically stratified (i.e., statistical areas), and the high-resolution data of real-time events (resident report data) and environmental factors. With this, it should be mentioned that the compensatory processes we observed are presumably density-dependent. Therefore, generalization of our results to areas with low cat densities should be taken cautiously. In addition, such a large study might also suffer from other limitations. For example, some processes might affect study results, such as changes in human behavior, changes in the urban environment (e.g., constructions), local changes in cat numbers, and even stochastic changes during the study period. We excluded the effect of these processes by controlling TNR both in space and time, incorporating possible confounders in the analysis, performing comparisons of the two datasets (resident data and annual counts data performed by experts), and comparing our data to datasets representing mortality of other animals and human behavior.

One limitation is a possible overestimation of the number of neutered cats due to a higher probability of detecting sociable cats than shy cats. These sociable cats are also more likely to be captured and neutered before capturing shy cats. Using friendly calling to increase the counted cat number might have enhanced this bias. However, a recent study showed that while friendly calling significantly increased the number of detected cats by 79%, its effect on the estimated neutering percentage was insignificant. Moreover, the mean insignificant increase by 18% (from 46% to 56%) of neutering percentage diminished almost completely when the neutering percentage was high (above 74%). This observation was probably due to the growing proportion of neutered shy cats as the neutering percentage increased (81). Therefore, despite the inherent uncertainty in the number of neutered cats not seen by the observers, we assume that our estimations of neutering intensity are reliable and valid and thus enable robust evidence for TNR effects and limitations.

### Conclusion

Our results indicate that fertility control can be applied to manage and even decrease the size of open free-roaming cat populations. However, the enhancement of compensatory mechanisms and the influx of immigrant cats might limit this effect. Therefore, we conclude that maintaining a high neutering rate for a prolonged period and in spatial contiguity is necessary to reduce cat population size. We recommend integrating these with complementary methods such as vital resource regulation, ill cat euthanasia, and adoption. In a broader view, our study suggests that the success of TNR should be measured by considering its environmental context, aims, and public acceptance. While simulation studies may aid in assessing such considerations, for enabling realistic interpretation, we recommend including parameters of possible compensatory effects and density-dependent immigration in these models. In particular, we call for a future robust evaluation of the effectiveness of integrated approaches.

## Materials and Methods

### Study site

We conducted the study from 2007 to 2018 in the city of Rishon-LeZion, located within the greater Tel-Aviv metropolitan area in Israel. The jurisdictional area of Rishon-LeZion is 58.7 km^2^, divided according to the Israeli Central Bureau of Statistics into 64 commercial and residential statistical areas. Three statistical areas spanning 33.8 Km^2^ include an isolated research institute, industrial and non-inhabited areas, and, therefore, not part of the study area. The city’s human population comprised 240,666 residents living in a 25 km^2^ area at the end of 2014 (Central Bureau of Statistics, Israel), divided into 58 residential areas (each consists of approximately 4000 residents) and three commercial areas (**Figure S1**). These 61 statistical areas constituted the study site.

### Study design

The study period was divided into three consecutive phases (**Figure 1, Table S1**): (1) the pre-intervention phase, (2) the mixed-intervention phase, and (3) the full-intervention phase. The pre-intervention phase preceded the initiation of the TNR program from January 2007 to the end of 2009. In the mixed-intervention phase, we allocated the 61 statistical areas into high-and low-intervention. In 31 statistical areas, the municipality implemented a high-intensity multi-annual TNR program from October 2009 until October 2014. The municipal TNR actions were limited in the other 30 statistical areas. We attempted to allocate TNR so that the human-related characteristics will be comparable between the high and low TNR regions but still maintain spatial contiguity among the un-neutered and neutered areas (see **Table S2** for demographic comparison between the high and low intervention areas). During this phase, 10,925 free-roaming cats were neutered with a male to female ratio of 1:1.07. In the full-intervention phase (November 2014 until December 2018), high-TNR efforts were expanded to the entire city. During this phase, 11,219 free-roaming cats were neutered with a male to female ratio of 1:1.06. The specific cat trapping and neutering procedures are detailed in the **supplementary information**.

### Data collection

To achieve high-resolution data on cat population dynamics, we used two independent data collection methods—a repeated annual survey and daily resident reports.

#### Annual surveys

We counted free-roaming cats via repeated annual surveys performed during September and October of 2012, 2013, 2014, and 2018. We conducted these surveys in 50 statistical areas (n=26 in the low-TNR statistical areas and n=24 in the high-TNR statistical areas) along fixed walking transects. In each statistical area, we chose one or two transects using a stratified random sampling design. Each statistical area was sampled by two observers, with each transect sampled twice each year. To increase the number of the observed cats the observer used friendly calling. We documented the individual characteristics of each observed cat, including sex, presumptive age (kitten<6 months vs. adults > 6 months), and sterilization status (according to the presence/absence of ear-marks). We summarized the annual counts of cats (adults (neutered and intact), intact-females (queens)), kittens (<6 months of age), and kitten-to-queen ratio per statistical area and year. We calculated the annual neutering percentage by dividing the total counts of neutered cats observed in each statistical area and year by the total counts of observed cats with identified neutering status. The transect walks were recorded using a cellular GPS recording application (“Endomondo™—Running & Walking, Android application,” Under Armour Inc., Maryland, USA), to ensure walking on precisely the same path of each transect and to measure the transect length. The total length of the transects was 100.87 km (**Figure S3**). A comprehensive description of this sampling method, including its intra-observer, inter-observer, inter-transect agreement, and validity, is detailed in Gunther et al. 2020 (81).

#### Resident reports

We retrieved data on cat carcasses and reproduction during 2007-2018 from the continuously available municipal emergency call center. The center receives voice complaints and reports from concerned residents on real-time events in the city jurisdiction area. The following data were recorded for each reported event: time and date of the call, location of the event, personal details about the calling resident, and a synopsis of the reported event. To confirm that the observed trends were unique to cat-related reports, we compared them to other municipal-based reports. The 2007-2018 cat carcass reports were compared to reports of carcasses of other animals (n=7,535), and cat reproduction reports were compared to the general municipal reports (apart from cat-related reports, n= 3,725,873) **(Figure S4**). Further information on the call center data and its retrieval methods are detailed in the **supplementary information**.

#### Environmental human-related data

The following environmental human-related factors were collected for Rishon-LeZion statistical areas during the study period: *human population density, socioeconomic status*, volume and type of *waste bins, cat feeding locations, educational institutions, food marketing businesses, built-up area* and *park area, age of neighborhood (years) and TNR actions performed by the municipality*. We provide further details on these factors in the **supplementary information**.

### Statistical analysis

Data were aggregated for each combination of statistical area and year. These values constituted the sampling units in all analyses. The following outcome variables were analyzed: cat counts, kitten counts, queen counts, kitten-to-queen ratio, neutering percentage, cat carcass reports, and cat reproduction reports. Except for the time-series analyses, all statistical analyses were performed by fitting generalized linear models (GLM) or generalized linear mixed models (GLMM) to these data (where the statistical area was used as a random effect). In all of these models, a negative binomial distribution was used. Since we analyzed rates, we used offsets as denominators (an offset is a variable that is forced to have a coefficient of 1 in the model. It is used in Poisson (or negative binomial) models to model rates. see **supplementary information for details)**. To account for potential spatial autocorrelation, we corrected the analyses by using an exponential variogram.

We performed the statistical analysis in two stages. The trends in the entire city were explored using all available data (i.e., survey data from 50 statistical areas and resident reports from 61 statistical areas). To discern the effect of TNR intensity on the cat population dynamics, we compared the cat population trends between the statistical areas with the highest to the lowest neutering intensity at the end of the mixed-intervention phase (designated as group-level analysis). To accomplish this, we divided the surveyed statistical areas into quartiles according to the observed neutering percentage at the end of the mixed-intervention phase. We compared two groups: Group-1 (n=12) consisted of the statistical areas with the lowest quartile of neutering percentage at this stage (≤30%), and Group-2 (n=15) consisted of the highest quartile of neutering percentage at this stage (≥75% with four statistical areas roughly tied at this threshold).

First, the effects of human-related factors (predictor variables) on the various cat population parameters in the entire city and their pairwise interactions with neutering were tested by GLM (outcome variables: surveyed cat and kitten counts, and cat carcass and reproduction reports). As we found that human population density and the volume of accessed waste bins per resident were the main human-related predictors of most cat population parameters, these were used in the following group-level analyses as covariates. To test the difference of cat population outcome variables between the annual surveys in the entire city, we performed GLMM. The group-level trend analyses were performed separately for the survey and resident report data. In both analyses, the effects of the treatment group, study phase, and their interaction (predictor variables) on the cat population parameters (outcome variables, as detailed above, plus queen counts and kitten-to queen ratio) were explored by GLMM. However, in the analysis of the resident report data, the year was added as a covariate in order to realize the high resolution of this data type. The specific modeling approach and statistical software used in each analysis are detailed in the **supplementary information**.

To visualize the temporal trends of the high-resolution resident report data (cat carcass and reproduction reports) collected both over the entire city or at the group level, we used time-series analysis (Seasonal and Trend decomposition using Loess (STL) method (82)). Through this method, we decomposed the monthly reports of the entire city (61statistical areas) into three compartments: the general trend, cyclic changes (designated as seasonality), and residual variability (remainder).

To confirm the construct validity of resident report data, we examined the correlation between reproduction reports and the survey-derived kitten counts. We aggregated the data to the neighborhood level and calculated the Pearson correlation coefficient of the log-transformed kitten counts + 0.5 per transect length and the log-transformed reproduction reports per the overall street length in each neighborhood in each year. (**Figure S6**).

## Acknowledgments

The authors are thankful to Y. Even-Zor, head of the municipal veterinary services, E. Gian, head of the municipal GIS department, S. Keidar, principal at the maintenance department, I. Ashkenazy from the municipal call center and E. Levi, head of the municipal information and research center at the city of Rishon-LeZion.

