## supplementary material for "Reduction of free-roaming cat population requires high-intensity neutering in spatial contiguity to mitigate compensatory effects"

**This PDF file includes:**

Supplementary text

Figures S1 to S6

Tables S1 to S4

Legend for Dataset_S1.xlsx

SI References

### Supplementary Information Text

#### Procedures

##### *Cat trapping*

Cats were located throughout the target neighborhoods using opportunistic trapping of individuals by municipal inspectors or in collaboration with cat caretakers. Thus, within the statistical areas allocated to high-intensity neutering, neutering was first performed in areas inhabited by large populations of un-neutered cats. Trapping was routinely performed using trigger-plate traps. In regions where neutering percentages reached high levels (approximately 70%), and to catch trap-shy cats, specific trapping procedures were selectively used: nets, traps that were triggered by remote control, shooting sedative drugs by blow-pipe, and accepting assistance from cat feeders (people who regularly fed cat colonies) that had gained the personal trust of specific cats.

##### *Neutering procedures*

Cats that underwent ovariohysterectomy or castration procedures were marked by cutting the tip of the left or right ear. The "ear-tipping" method visually signals that a cat has been sterilized (i.e., spayed/neutered) and vaccinated against rabies. Marking was performed under general anesthesia during the sterilization procedure (1). Following recovery, cats were returned to the same location where they had been trapped. Municipal Veterinary Services kept meticulous records for each neutered cat, including the trapping date and location (documented as the street address closest to the trapping location).

#### Data

##### *Description of the call center data and their retrieval*

We retrieved resident reports regarding *cat carcasses* from the general animal-related reports using a simple keyword algorithm containing the word “cat”. Non-identified carcasses and duplicate reports were omitted. The remaining reports were geographically coded and summarized by statistical area. Resident reports regarding *free-roaming cat reproduction* were those reports that contained the words “cat” and one of the following words: “to give birth”; “to be born”; “pregnant”; “kitten”. This was performed by using a computerized specific keyword algorithm. Further data-cleaning of duplicate reports and geographical coding was performed similarly to the carcass reports. For a more comprehensive explanation of the data-mining process, see Gunther et al. 2015 (2). Overall, 3,759,530 reports were recorded in the call center during the study period (2007-2018) in the entire city (61 statistical areas). Out of these reports, 12,217 were on cat reproduction and 44,079 were identified animal carcass reports, from which 36,544 cat carcass reports were identified and analyzed.

***Environmental human-related data***

1. The *number of residents* in 2009, 2014 and 2017 was determined by the Central Bureau of Statistics, Israel. Data were divided by the area of each statistical area in $km^2$ and were used as the *human population density*.
2. *Socioeconomic status* in 2008 was determined by the Central Bureau of Statistics, Israel, based on the 2008 national census. The estimated socioeconomic status was measured on a continuous scale ranging from 389 to 1395 (a higher number represents higher socioeconomic status). These ranks were calculated by the Israeli Central Bureau of Statistics and represent a combination of variables including demographic composition, education, labor, housing, and income.
3. Data of *waste bins* was documented by the Municipal Maintenance Department during the end of 2012 and 2018. Waste bin data consist of the location, volume, and bin type (e.g., closed bins, closed and open dumpsters, underground waste containers, and garbage compactors). Data were divided into three sub-categories according to the potential accessibility of waste to cats: inaccessible (underground waste containers and garbage compactors), partially accessible (closed bins and dumpsters), and fully accessible (open dumpsters). In addition to testing each sub-category separately, the partially and fully accessible waste bins were combined and modeled together. Data were geographically coded, summarized as total waste-bin-volume-per-statistical-area, and then standardized by the number of residents.
4. *Cat feeding locations* were reported in the 2013 telephone survey conducted among cat feeders (for more details, see Gunther et al. 2016 (3)). The locations were geographically coded, summarized per statistical area, and standardized by the number

of residents. Due to the length of the study period, this factor was excluded from the 2018 analyses.

5. The location, number, and type of *educational institutions* were documented for 2014 and 2017 by the Municipal GIS Department. Educational institutions were summarized per statistical area and standardized by the number of residents.
6. The type and location of *food service businesses* (e.g., butcher shops, restaurants, supermarkets, catering) were documented for 2012 and 2018 by the Municipal Department for Business Registrations and Municipal Veterinary Services. *Food service businesses* were geographically coded, summarized per statistical area, and standardized by the number of residents.
7. The *built-up area* and *park area* were documented for 2014 and 2017 by the Municipal GIS Department. Data were summarized per statistical area and standardized by its area in $Km^2$.
8. The *age of each neighborhood* was documented by the Municipal Information and Research Center. This information was relevant to the study as sanitation and potential hiding places available to cats depend on architecture and infrastructure, which differ substantially between neighborhoods according to their year of development. The oldest and youngest neighborhoods were established in 1882 and 2008, respectively.
9. *TNR actions* were geographically coded and summarized per statistical area and month. Three hundred forty-five trapped cats were omitted due to missing or unclear capture location information. Further, 3479 cat-trap sites were randomly chosen from a specific reporting area, such as parks or blocks.

### Statistical analysis

#### *Rate modelling using offsets*

To model rates, offsets where introduced in the models as follows. The counts of reproduction and carcass reports for each statistical area and year where modeled using the log-transformed 'Street length' (the overall length of street in each statistical area). The cat kitten and queen counts for each statistical area and year combination where modeled using the log-transformed 'Transect length' as an offset. The counts of kittens were modeled as a kitten-to-queen ratio using the log-transformed number of queens as an offset, and finally, the counts of neutered cats were modeled using the log-transformed ‘total count of cats’ as an offset.

***Specific modeling approach***

*Analysis of environmental human-related factors at city scale:*

The following model was used to screen for important human-related factors affecting cat population parameters (**Table S3**):

The following model was used to screen for important human-related factors affecting cat population parameters (**Table S3**):

*Equation 1:* $$Y_j = \beta_0 + \sum_{l=1}^{k}(\beta_l C_{l,j,i}) + \ln\left(F_{j,i}\right) + e_j$$

Where $Y_j$ is the log transformed outcome (count variable) for the $j^{th}$ statistical area, $C_{l,j,i}$ is the $l^{th}$ covariate for the $j^{th}$ statistical area in the $i^{th}$ year (see **Table S2** for a detailed list of all human related covariates), $F_{j,i}$ is the offset for the $j^{th}$ statistical area in the $i^{th}$ year, and $e_j$ is the residual. Analyses were performed for each year separately (i.e., for years 2012–2014 and 2018). Owing to the numerous human-related variables, we performed the analysis in five steps. First, we conducted a univariable analysis separately for each covariate. Second, we included strong covariates ($p<0.05$) in a multivariable analysis after ruling out collinearity. Third, we included interactions between each covariate and the annual estimated neutering percentage. Forth, using 'model average analysis,' the most parsimonious model was selected using the Akaike-Information-Criterion (AIC). A model with an increased number of covariates was selected if its AIC was smaller by at least three units. Finally, to account for potential spatial autocorrelation, we corrected the model by using an exponential variogram.

*Survey results: Analysis of the effect of year at city scale:*

The following model was used to analyze the differences between years 2012, 2013, 2014 and 2018 in cat counts, kitten counts, queen counts, kitten-to-queen ratio and neutering percentage in the entire city:

*Equation 2:* $$Y_j = \beta_0 + \beta_{1i}X_i + \ln(F_{j,i}) + v_{0j,i} + e_j$$

Where $Y_j$ is the log transformed outcome (count variable) for the $j^{th}$ statistical area, $X_i$ is an indicator of the $i^{th}$ year effect (i.e. 2012, 2013, 2014 or 2018), $F_{j,i}$ is the offset for the $j^{th}$ statistical area in the $i^{th}$ year, $v_{0j,i}$ is the random effect on the intercept for the $j^{th}$ statistical area in the $i^{th}$ year and $e_j$ is the residual.

*Survey results: Analysis of trends across treatment groups and study phases*

The following model was used to analyze the trends of cat, kitten and queen counts and kitten-to-queen ratios per treatment group and study phase:

*Equation 3:* $$Y_j = \beta_0 + \beta_{1i}X_i + \beta_{2k}G_k + \beta_{3i,k}X_iG_k + \beta_4 P_{j,l} + \beta_5 W_{j,l} + \ln(F_{j,i}) + v_{0j,i} + e_j$$

Where $Y_j$ is the log transformed outcome (count variable) for the $j^{th}$ statistical area, $X_i$ is an indicator of the $i^{th}$ year effect (i.e. 2012, 2014 or 2018), $G_k$ is the $k^{th}$ group effect indicator (i.e. Group-1 or Group-2), $P_{j,l}$ is the human population density as a covariate for the $j^{th}$ statistical area at the $l^{th}$ phase, $W_{j,l}$ is the volume of accessed waste bins per resident as a covariate for the $j^{th}$ statistical area at the $l^{th}$ phase, $F_{j,i}$ is the offset for the $j^{th}$ statistical area in the $i^{th}$ year, $v_{0j,i}$ is the random effect on the intercept for the $j^{th}$ statistical area in the $i^{th}$ year and $e_j$ is the residual.

The model results were interpreted as follows: $\beta_{1i}$ is the difference between two years for a specific group, thus representing the group specific phase trend (2012-2014 – mixed-intervention phase, 2014-2018 – full intervention phase). $\beta_{3i,k}$ is the trend difference between the two groups within a specific phase. For interpretation, the estimated regression coefficients were exponentiated, and for an interpretation at annual level, the estimated parameters were weighted by the number of years of the respective phases (i.e. weighted by 2 years for the mixed intervention phase, and 4 years for the full-intervention phase). The resulting trend estimates were presented in percent. The statistical differences between the trends of the two phases within each group were calculated using the ‘comparing two ratios’ module in WinPepi (4).

*Resident report results: Analysis of trends across treatment groups and study phases*

following model was used to assess the trends of cat carcass and reproduction reports across treatment groups and study phases:

*Equation 4:*

$$Y_j = \beta_0 + \beta_{1l}X_l + \beta_{2k}G_k + \beta_{3l}Z_l + \beta_{4k}XG_k + \beta_{5l}XZ_l + \beta_{6l,k}Z_lG_k + \beta_{7l,k}XZ_lG_k + \beta_8P_{j,l} + \beta_9W_{j,l} + ln(F_{j,i}) + v_{0j,i} + e_j$$

Where $Y_j$ is the log transformed outcome (count variable) for the $j^{th}$ statistical area, $X_l$ is the year as a covariate in $l^{th}$ phase, $G_k$ is the $k^{th}$ group effect indicator (i.e. Group-1 or Group-2), $Z_l$ is an indicator of $l^{th}$ phase effect (i.e., 1st phase - pre-intervention period (2007-2009), 2nd phase – mixed-intervention period [2010-2014], and 3rd phase – full-intervention period [2015-2018]), $P_{j,l}$ is the human population density as a covariate for the $j^{th}$ statistical area in the *lth* phase, $W_{j,l}$ is the volume of accessed-waste-bins-per-resident as a covariate for the $j^{th}$ statistical area in the $l^{th}$ phase, $v_{0j,i}$ is the random effect on the intercept for the $j^{th}$ statistical area in the $i^{th}$ ear and $e_j$ is the residual. Note that the resident report data were available for every year from 2007 to 2018. Therefore, as opposed to the previous model (*equation 3)*, the year was modeled as a covariate to take advantage of the higher resolution of data.

The model results were interpreted as follows: $\beta_{1l}$ is the average annual trend for a specific group during a specific phase. $\beta_{4k}$ indicates the trend difference between the groups during a specific phase. $\beta_{5l}$ indicates the trend difference between two phases within a specific group. For interpretation, the estimated regression coefficients were exponentiated. The resulting trend estimates were presented in percent.

***Software***

Geographical data were presented on ArcGIS Desktop version 10.5.1 (ESRI 2011. Redlands, CA: Environmental Systems Research Institute). We performed the statistical analyses using the following packages in R (5): 'nlme' and 'lme4' for generalized linear models (GLM) and for generalized linear mixed models (GLMM) (6, 7); 'MuMIn' for model averaging analysis (8); 'forecast' for time series analysis; 'broom' and 'ggplot2' for model diagnostic analysis and generating figures, respectively (9, 10); ‘MASS’ for spatial auto-correlation analysis; and ‘car’ for model diagnostics analysis (11, 12). In all analyses, a 5% significance level α was applied.

**Figures:**

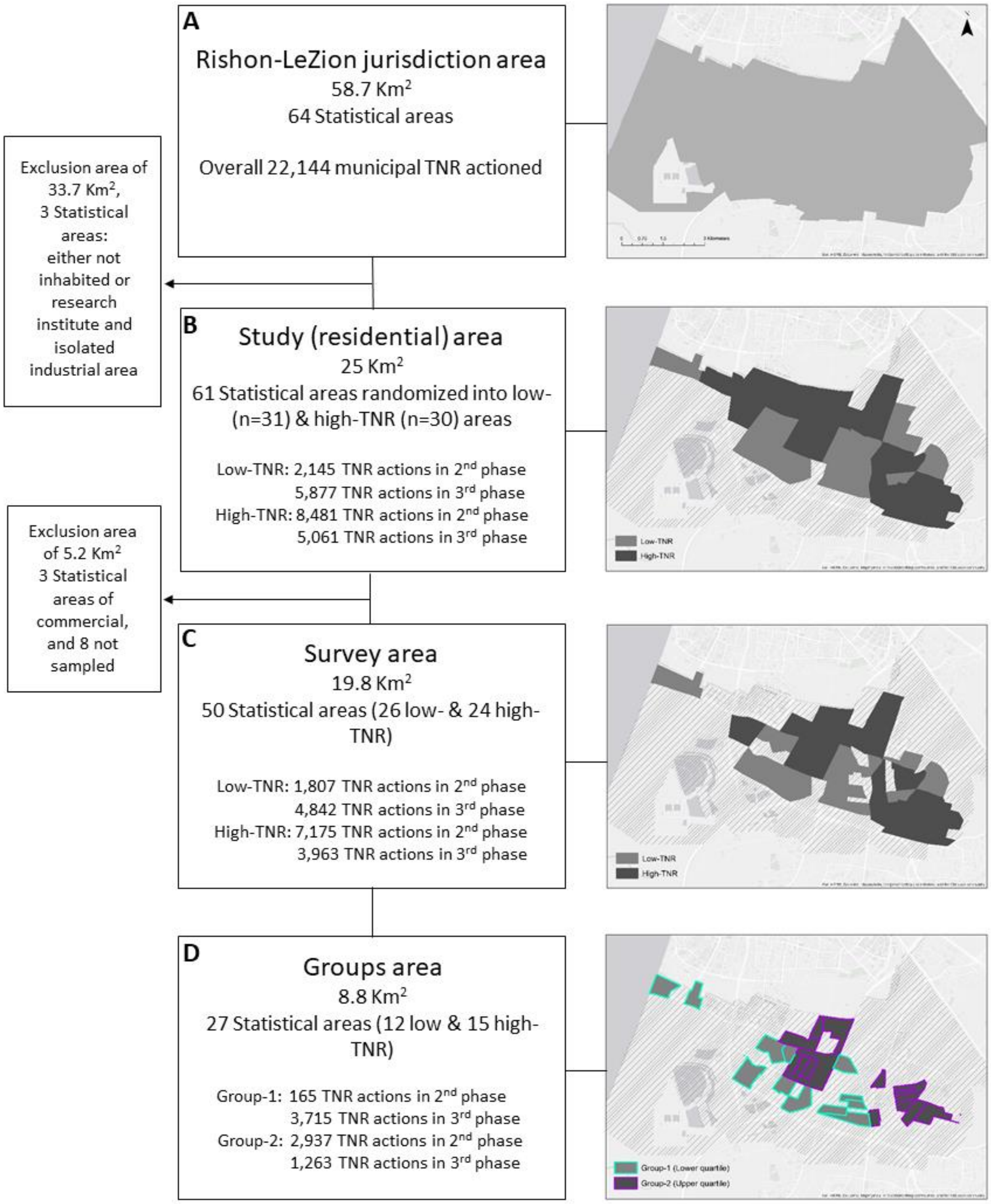


**Figure S1**: Spatial design of the study: **A**. The entire jurisdiction area of Rishon-Lezion. **B.** Allocation of TNR (dark grey) to 30 of the 61 statistical areas in the study area during the mixed-intervention period. These areas were designated as high-TNR areas. **C.** Survey area of free-roaming cats in 50 statistical areas. **D.** Sub-division into low- (Group-1, turquoise edges) and high-quartile (Group-2, purple edges) according to the observed neutering percentage at the end of 2014 (end of the 2nd phase). Arrows pointing outside indicate unsampled area.

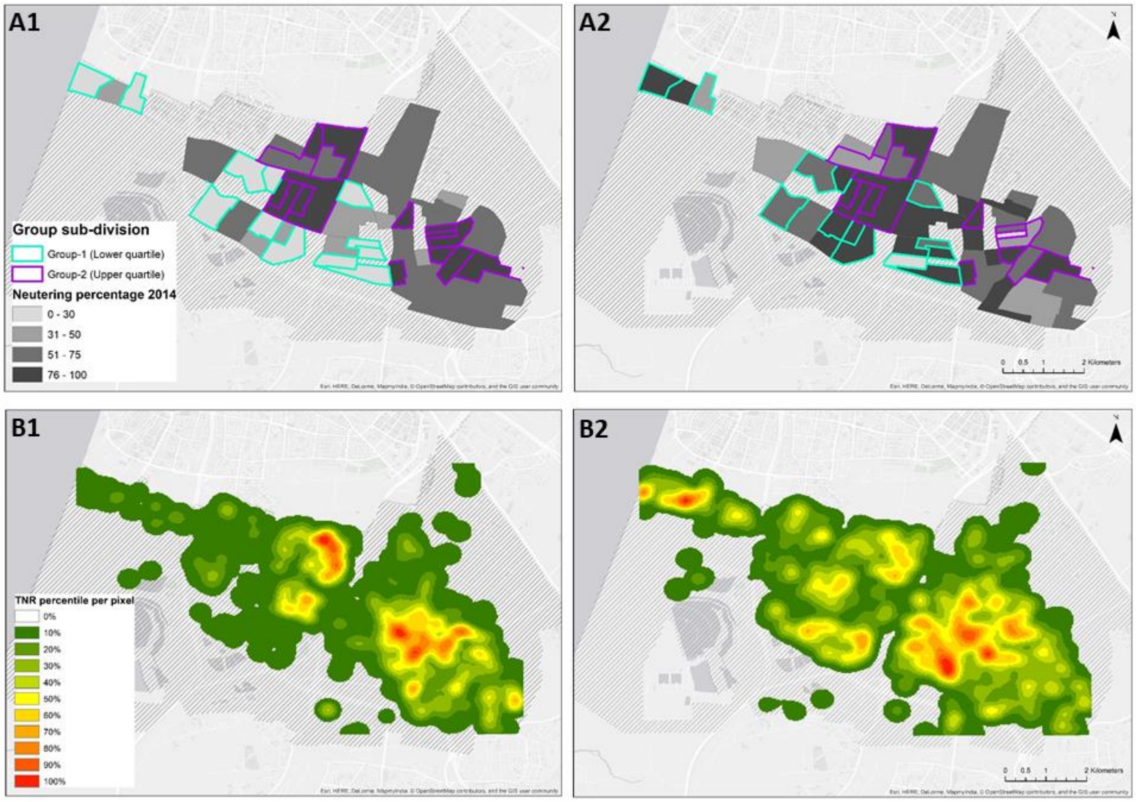


**Figure S2**: Spatial distribution of the observed neutering percentages in the surveyed statistical areas (n=50)(**A**) and the corresponding heat maps of municipal TNR efforts in the entire city (n=61 statistical areas) (**B**), at the end of the mixed-intervention phase (2014) (**A1,B1**) and the full-intervention phase (2018) (**A2,B2**). Group-1 (turquoise edges) and Group-2 (purple edges), include the statistical areas that received the lowest and highest TNR intensity, respectively.

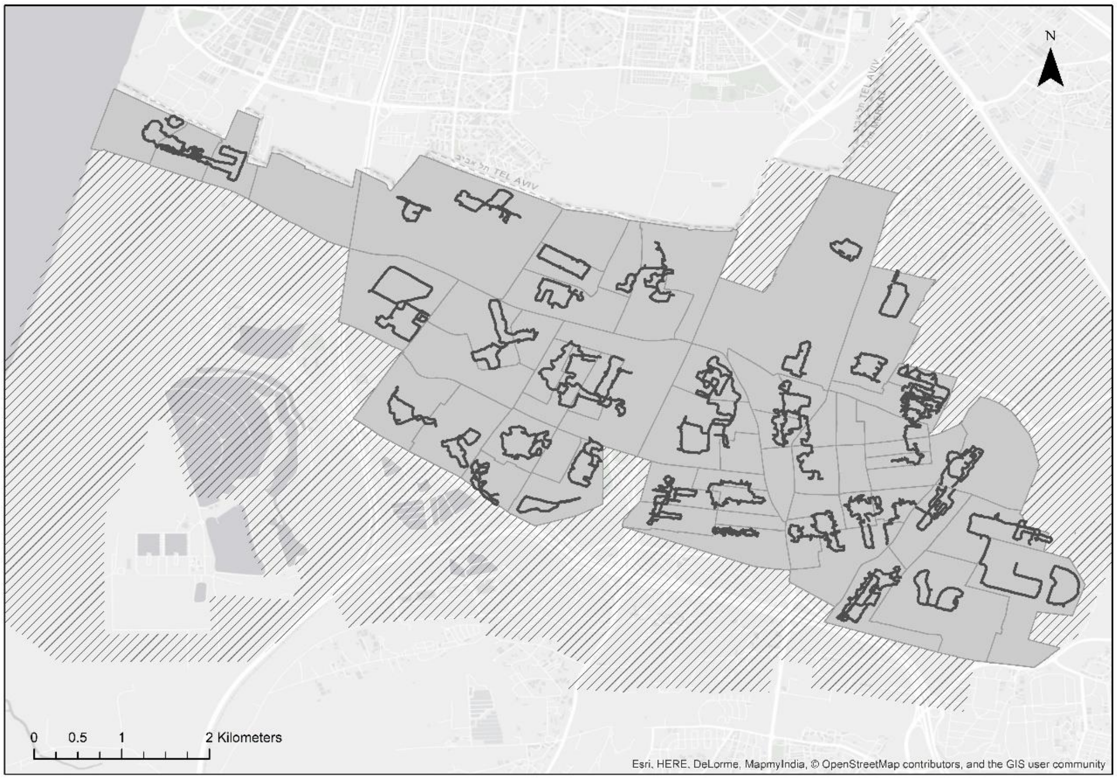


**Figure S3:** A map of the observation transects walked during the study. The survey design included randomly chosen transects (overall length of 100.87 km) in 50 statistical areas of the city of Rishon-LeZion. Observations were performed twice in each year (2012, 2013, 2014, 2018).

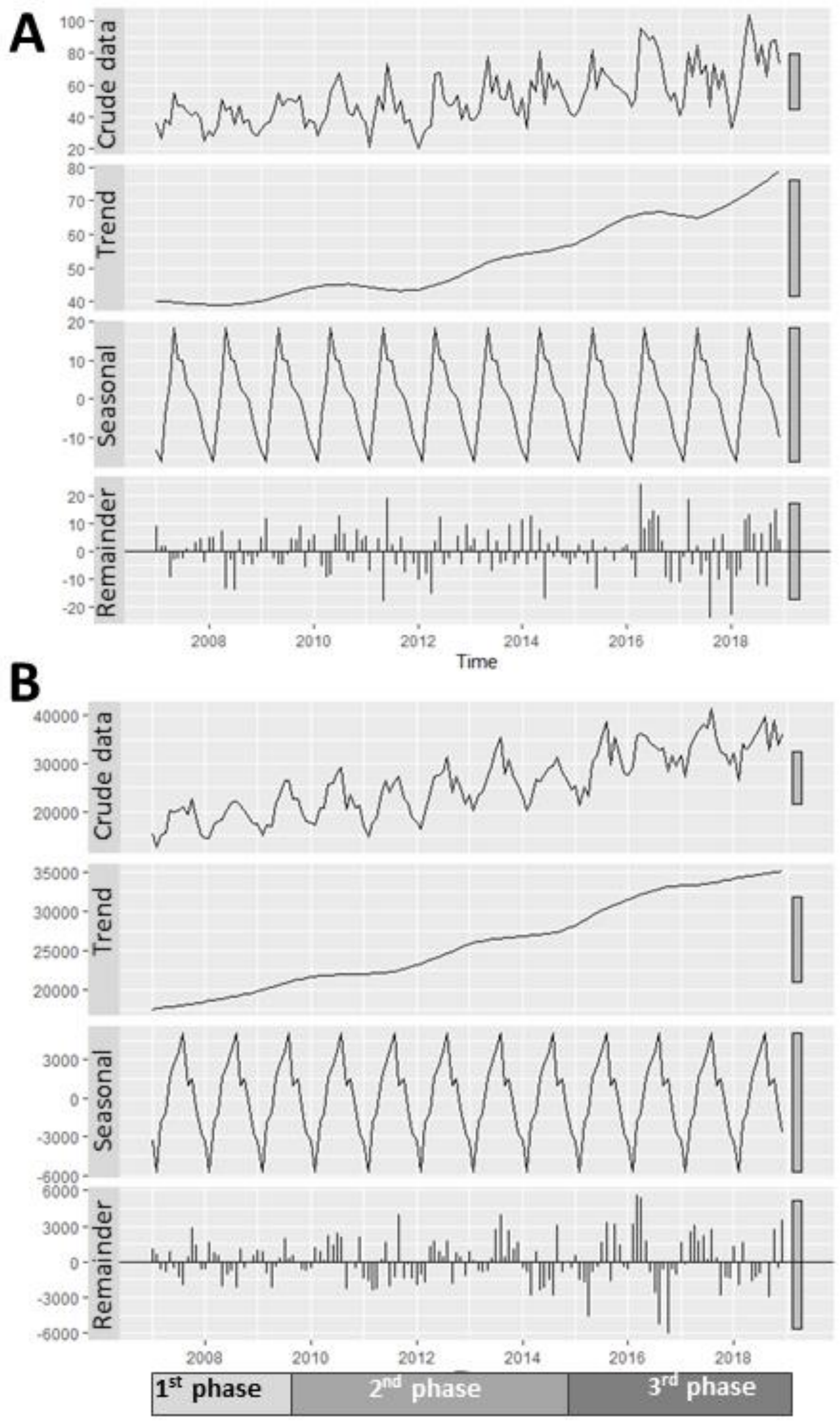


**Figure S4**: Monthly-based time series decomposition plot (between 2007-2018; the crude data is divided into three compartments: trend, seasonality and remainder) of (**A**) animal carcasses (canines, poultry, rodents, hyrax, hedgehogs, equines, reptiles, boars, tortoises) (n=7,535), and (**B**) general resident reports (n= 3,725,873). Both reports types were registered by the municipal call-center from January 2007 to December 2018 across the entire city and were reported after excluding cat records.

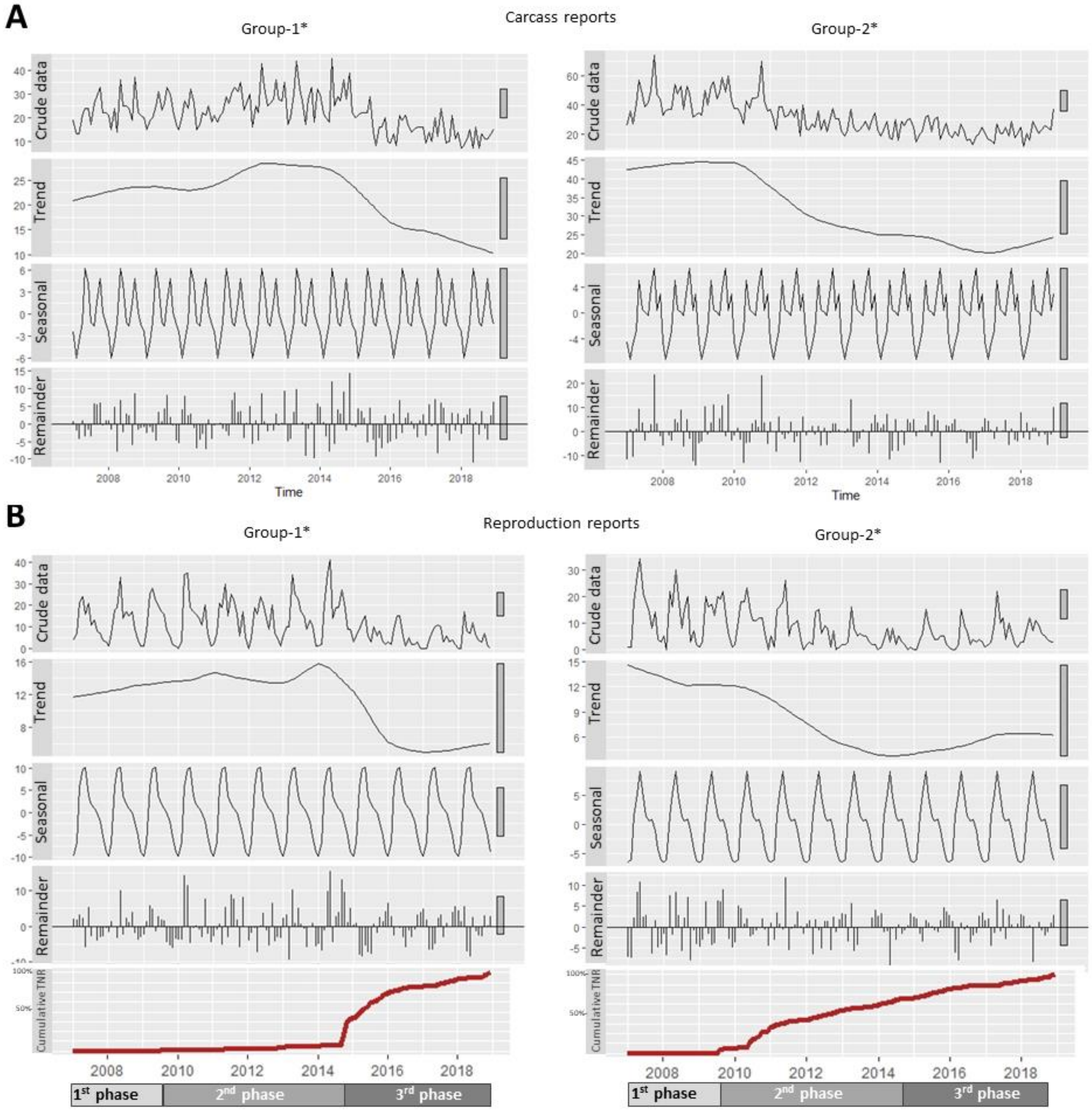


**Figure S5**: Cumulative TNR percentage and monthly-based time series decomposition plots (between 2007-2018; the crude data is divided into three compartments: trend, seasonality and remainder) of (**A**) cat carcass reports and (**B**) cat reproduction reports in Group-1* statistical areas (n = 7) compared to Group-2* statistical areas (n = 9), in which the observed neutering percentage in 2018 was above 70%.

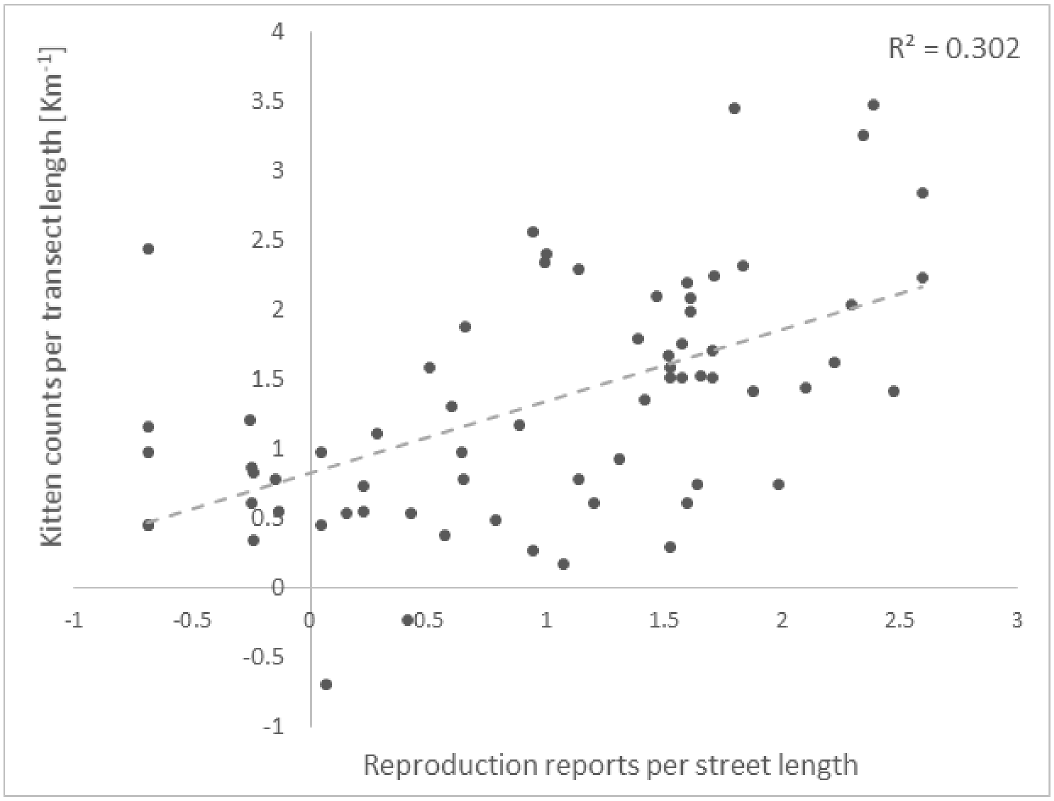


**Figure S6:** Scatterplot of the year and neighborhood specific log-transformed kitten counts per street length against the log-transformed reproduction reports per street length. (Pearson correlation coefficient = 0.55, $p<0.001$).

**Table S1**. TNR efforts by the municipality in the mixed-intervention phase (2nd phase, between the end of 2009 to the end of 2014), and in the full-intervention phase (3rd phase, between the end of 2014 to 2018). Low-TNR and high-TNR statistical areas are those that were randomized during the 2nd phase to low and high intensity TNR actions, respectively. From these areas, Group-1 and Group-2 represent the areas with the lowest and highest neutering percentage at the end of the 2nd phase (end of 2014).

| Group division (number of statistical areas) | Low-TNR (n=26) | High-TNR (n=24) | Group-1 (n=12) | Group-2 (n=15) |
|---|---|---|---|---|
| Observed cats in 2012 per km transect | 16.99 | 18.23 | 16.01 | 23.35 |
| Street length [Km] | 134.71 | 180.26 | 56.88 | 63.95 |
| Proxy cat abundance in 2012* | 2288.72 | 3286.14 | 910.65 | 1493.23 |
| Annual TNR actions in 2nd phase: | | | | |
| 2009 | 38 | 393 | 23 | 326 |
| 2010 | 391 | 1908 | 12 | 1178 |
| 2011 | 662 | 1410 | 18 | 764 |
| 2012 | 362 | 1447 | 43 | 609 |
| 2013 | 205 | 980 | 49 | 397 |
| 2014 up to Sep. | 149 | 1013 | 20 | 441 |
| 2nd phase overall TNR actions divided by proxy cat abundance in 2012 | 0.72 | 1.9 | 0.18 | 2.48 |
| Annual TNR actions in 3rd phase: | | | | |
| 2014 Oct. to Dec | 1509 | 130 | 841 | 50 |
| 2015 | 1711 | 1559 | 806 | 602 |
| 2016 | 615 | 788 | 236 | 278 |
| 2017 | 440 | 535 | 213 | 283 |
| 2018 | 567 | 951 | 256 | 482 |
| 3rd phase overall TNR actions divided by proxy cat abundance in 2012 | 2.12 | 1.2 | 3 | 0.95 |
| Overall TNR actions | 6649 | 11114 | 2517 | 5410 |
| Overall TNR actions divided by proxy cat abundance in 2012 | 2.9 | 3.38 | 2.76 | 3.62 |

* We calculated cat-adjusted TNR intensity by dividing the number of TNR actions (municipality neutered cats) by a proxy abundance of free-roaming cats per each group in 2012. The proxy cat abundance was calculated by dividing the survey cat counts in 2012 per transect length and multiplying it by the total street length of each area.

**Table S2**. Environmental human-related characteristics of the low TNR statistical areas (n=26), high TNR statistical areas (n=24), Group-1 (lower quartile of the distribution of the observed neutering percentage in 2014, n=12) and Group-2 (higher quartile, n=15).

| Characteristics | Low-TNR | High-TNR | Group-1 | Group-2 |
|---|---|---|---|---|
| Overall area [$Km^2$] | 7.95 | 11.82 | 4.08 | 4.71 |
| Human population [mean±SD] | 3980 ± 1382 | 4408 ± 1260 | 3665 ± 1466 | 4101 ± 1048 |
| Socio-economic status [mean±SD] | 1174± 141 | 945 ± 326 | 1211 ± 116 | 974 ± 331 |
| Feeding locations per 1000 residents [mean±SD] | 1.28 ± 0.97 | 1.51 ± 1.01 | 1.41 ± 0.79 | 1.25 ± 0.99 |
| Accessed waste bins [mean±SD] | 36.2 ± 9.5 | 38.6 ± 10.4 | 36.75 ± 9.3 | 33.55 ± 7.17 |
| Educational institutes per 1000 residents [mean±SD] | 1.62 ± 0.90 | 1.56 ± 0.8 | 1.90 ± 1.10 | 1.66 ± 2.12 |
| Food businesses per 1000 residents [mean±SD] | 0.44 ± 0.46 | 0.52 ± 0.71 | 0.27 ± 0.33 | 0.23 ±0.42 |
| Neighborhood age (years) [mean±SD] | 49.46 ± 33.19 | 68.17 ± 43.58 | 39.33 ± 31.86 | 72.93 ±44.48 |
| Area of parks [mean±SD] | 0.11 ± 0.16 | 0.05 ± 0.15 | 0.13 ±0.15 | 0.06 ± 0.19 |
| Built-up area [mean±SD] | 0.22 ± 0.07 | 0.24 ± 0.08 | 0.20 ±0.08 | 0.25 ± 0.1 |

**Table S3**: Environmental human-related factors associated with the annual cat population parameters between 2012─2014 (mixed-intervention phase) and in 2018 (end of full-intervention phase). The rate ratio per factor unit and the distance of spatial correlation are presented for each variable in each year.

| Outcome variable | Year | Anthropogenic/ environmental factor | Rate ratio ($CI_{95\%}$) | Distance of spatial correlation [m] |
|---|---|---|---|---|
| Cat counts | 2012 | Measured neutering proportion* | 1.068 (1.013 to 1.126) | Ng |
| | | Human population density** | 1.226 (1.041 to 1.443) | |
| | 2013 | Human population density** | 1.342 (1.143 to 1.575) | NA |
| | 2014 | Full accessed waste bins£ | 1.042 (1.005 to 1.08) | NA |
| | 2018 | Measured neutering proportion * | 1.062 (0.979 to 1.152) | Ng |
| | | Human population density** | 1.164 (0.957 to 1.415) | |
| Kitten counts | 2012 | NA | NA | NA |
| | 2013 | Measured neutering proportion * | 0.850 (0.771 to 0.936) | Ng |
| | 2014 | Measured neutering proportion * | 0.827 (0.751 to 0.910) | 191 |
| | 2018 | Measured neutering proportion * | 0.835 (0.719 to 0.967) | Ng |
| | | Educational institutes** | 0.961 (0.929 to 0.993) | |
| Carcass reports | 2012 | Human population density** | 1.294 (1.122 to 1.495) | Ng |
| | | Area of parks ($Km^2$) | 0.359 (0.173 to 0.743) | |
| | 2013 | Human population density** | 1.505 (1.331 to 1.702) | Ng |
| | | Zero accessed waste bins | 0.981 (0.966 to 0.996) | |
| | 2014 | Measured neutering proportion * | 0.906 (0.873 to 0.940) | Ng |
| | | Human population density** | 1.859 (1.476 to 2.342) | |
| | | Neighborhood seniority (years) | 1.012 (1.006 to 1.018) | |
| | | Interaction of human population density and Neighborhood seniority | 0.996 (0.993 to 0.999) | |
| | 2018 | Full and partial accessed waste bins£ | 1.020 (1.012 to 1.028) | 220 |
| Reproduction reports | 2012 | Measured neutering proportion * | 0.836 (0.784 to 0.896) | 74 |
| | | Human population density** | 1.610 (1.300 to 1.996) | |
| | 2013 | Measured neutering proportion * | 0.866 (0.796 to 0.942) | Ng |
| | 2014 | Measured neutering proportion * | 0.778 (0.733 to 0.825) | 79 |
| | | Human population density** | 1.584 (1.326 to 1.895) | |
| | 2018 | Measured neutering proportion * | 0.894 (0.826 to 0.969) | 214 |
| | | Full and partial accessed waste bins£ | 1.017 (1.006 to 1.027) | |

Ng = Negligible distance (less than 50 m) NA = Not Applicable
* Fits a theoretical 10% increase in neutering ** Presented per 10,000 residents.
£ Presented for a change of one liter per person

**Table S4:** Same as table 1 but including only statistical areas in which >70% neutering percentage was observed in 2018. The annual trend of FRC, kitten, queen counts in the mixed-intervention (2$^{nd}$) and full-intervention (3$^{rd}$) phases of the study: (7 statistical areas in Group-1* and 9 statistical areas in Group-2*).

| Variable | Phase | Measurement period | Annual trend for Group-1* ($CI_{95\%}$) [%] | Annual trend for Group-2* ($CI_{95\%}$) [%] | P-value |
|---|---|---|---|---|---|
| Cat counts | 2nd phase | 2012-2014 | 17.01 (1.28 to 35.19) [a1*] | 1.61 (-10.08 to 14.80) [a2] | 0.143 |
| | 3rd phase | 2014-2018 | -6.33 (-13.19 to 1.08) [b1] | -6.56 (-12.17 to -0.66) [a2] | 0.959 |
| Kitten counts | 2nd phase | 2012-2014 | 17.67 (-16.80 to 66.44) [a1] | -40.0 (-57.63 to -15.00) [a2] | 0.007 |
| | 3rd phase | 2014-2018 | -29.16 (-42.12 to -13.29)[b1] | 12.01 (-6.33 to 33.93) [b2] | <0.001 |
| Queen counts | 2nd phase | 2012-2014 | 37.69 (8.72 to 74.37) [a1] | -23.84 (-42.39 to 0.7) [a2] | 0.002 |
| | 3rd phase | 2014-2018 | -46.06 (-55.63 to -34.42) [b1] | -16.36 (-29.77 to -0.39) [a2] | <0.001 |
| Kitten-to-queen ratio | 2nd phase | 2012-2014 | -5.36 (-41.24 to 52.44) [a1] | -11.10 (-42.89 to 38.36) [a2] | 0.853 |
| | 3rd phase | 2014-2018 | 28.22 (-1.03 to 66.12)[a1] | 21.84 (-3.26 to 53.45) [a2] | 0.773 |
| Carcass reports | 1st phase | 2007-2009 | 4.58 (-7.47 to 18.20)[a1] | 3.81 (-5.88 to 14.49) [a2] | 0.926 |
| | 2nd phase | 2010-2014 | 5.27 (-0.22 to 11.07) [a1] | -12.46 (-16.54 to -8.17) [b2] | <0.001 |
| | 3rd phase | 2015-2018 | -13.17 (-20.43 to -5.25) [b1] | -2.21 (-9.11 to 5.21) [a2] | 0.041 |
| Reproduction reports | 1st phase | 2007-2009 | 4.24 (-15.27 to 28.25) [a1,b1] | -6.98 (-22.20 to 11.23) [a2] | 0.415 |
| | 2nd phase | 2010-2014 | 12.76 (3.34 to 23.04) [a1] | -28.54 (-35.22 to -21.16) [b2] | <0.001 |
| | 3rd phase | 2015-2018 | -11.45 (-23.72 to 2.80) [b1] | 10.68 (-3.91 to 27.48) [a2] | 0.033 |

* The superscript letters represent comparisons between pairs of phases within each group of SAs (pairs with insufficient evidence for a difference are labeled with the same superscript letter).

**Additional data:**

Dataset_S1.xlsx: An excel file including the data and code for the tables and figures in the main text and the supplementary information.
